# Macrophage cell lines and murine infection by *Salmonella* Typhi L-form bacteria

**DOI:** 10.1101/2021.12.10.472194

**Authors:** Debayan Ganguli, Swarnali Chakraborty, Suparna Chakraborty, Ananda Pal, Animesh Gope, Santasabuj Das

## Abstract

Antibiotic resistance of pathogenic bacteria has emerged as a major threat to public health worldwide. While stable resistance due to the acquisition of genomic mutations or plasmids carrying antibiotic-resistance genes is well-established, much less is known about the temporary and reversible resistance induced by antibiotic treatment, such as the one due to treatment with bacterial cell-wall inhibiting antibiotics like ampicillin. Typically, ampicillin concentration in the blood and other tissues gradually increases over time after initiation of the treatment. As a result, the bacterial population is exposed to a concentration gradient of ampicillin. This is different from *in vitro* drug testing where the organism is exposed to fixed drug concentrations from the beginning till the end. To mimic the mode of antibiotic exposure of microorganisms in the tissues, we cultured the wild type, ampicillin-sensitive *Salmonella* Typhi Ty2 strain (*S*. Typhi Ty2) in the presence of increasing concentrations of ampicillin over a period of 14 days. This resulted in the development of a strain that exhibited several features of the so-called L-form of bacteria, such as the absence of cell wall, altered shape and slower growth rate compared with the parental strain. Studies on the pathogenesis of *S*. Typhi L-form showed efficient infection of the murine and human macrophage cell lines. More importantly, *S*. Typhi L-form was also able to establish infection in a mouse model to the extent comparable to its parental strain. These results suggested that L-form generation following initiation of antibiotic treatment could lead to drug escape of *S*. Typhi and direct spread to new cells (macrophages), which sustain the infection. Oral infection by the L-form bacteria underscores the potential of rapid disease transmission through faeco-oral route, highlighting the need for new approaches to decrease the reservoir of infection.

## Introduction

Around 26.9 million cases of typhoid fever are reported annually worldwide [1]. Multi drug resistant strains of *Salmonella* Typhi are known for several decades and were isolated at different parts of the globe. *S*. Typhi was earlier treated with Ampicillin, chloramphenicol and trimethoprim–sulfamethoxazole [2]. Appearance of multidrug resistance to the first line antibiotics during late 1980s led fluoroquinolones, especially ciprofloxacin to become the drug of choice for typhoid fever [3]. In 1991, *S*. Typhi strain resistant to ciprofloxacin emerged, which lead to an outbreak in 1997 [4]. Subsequently, macrolides, 3^rd^ generation cephalosporins and carbapenems were widely used to treat *S*. Typhi infection. In 2018 and 2019, extreme drug resistant (XDR) *S*. Typhi strains, unresponsive to macrolides, fluoroquinolones and 3^rd^ generation of cephalosporins were reported from Hyderabad [5, 6], India and Pakistan. WHO had recently announced 10,365 infections with XDR strains in Pakistan and travel-associated infections in China, USA, Australia, Denmark and Canada, thus making *S*. Typhi drug resistance a global threat [7]. Interestingly, reversal to ampicillin sensitivity of *S*. Typhi was reported by studies carried out at different parts of India [8]. Harish *et al*, 2011 had reported that ampicillin-resistant strains in Chennai fell from 52% to 23% during the period 2002 to 2009 [9]. In another study, around 94.5 % of *S*. Typhi clinical isolates collected from India were found to be ampicillin sensitive [10].

Antibiotic resistance of *S*. Typhi was believed to be plasmid borne. Genome sequencing of CT18 and several other Ampicillin resistant *S*. Typhi strains revealed the presence of *Salmonella*-associated transferable R plasmid, called pHCM1 [11]. This plasmid was responsible for the resistance to all three first-line antibiotics. However, temporary and reversible ampicillin resistance may also be conferred by the formation of L-form bacteria. Ampicillin targets the bacterial cell wall machinery and bacteria can overcome the lethal effects of the drug by shutting down the cell wall production temporarily. Bacterial L-form was first discovered in 1935 [12] and subsequent studies characterized this form. However, most studies were carried out in the pre-molecular biology era and thorough knowledge about the formation of the L form is lacking. Recently, Errington and others explored the mechanism underlying L-form transition of *B. subtilis* [13, 14, 15]. They discovered that deletion of the cell wall machinery genes (MurE operon) was not sufficient to induce stable L-form growth. Inactivating mutation of ispA or a glycolytic or respiratory chain enzyme in addition to MurE knock out was necessary for the L-form to survive [15]. However, the transient and reversible nature of antibiotic resistance associated with the L-form failed to attract much global attention, leading to poor understanding about its significance in pathogenesis and contribution to treatment failures associated with antimicrobial resistance. Notwithstanding, bacterial L-forms were shown to be responsible for recurrent and chronic infections, although their diagnosis as etiologic agents is frequently missed due to slow growth and the requirement for specialized culture medium. Back in 1965, L-form was suggested to cause chronic Staphylococcal infection [16]. Gutman et al. isolated L-forms of Escherichia coli, Klebsiella spp. and Enterococcus faecalis from patients suffering from chronic bacteriuria or pyelonephritis [17]. *E. facealis* L-forms persisted in rats infected with walled bacteria, followed by treatment with penicillin [18]. Cell wall deficient L-forms of *Corynaebacterium* and *Mycobacterium tuberculosis* were isolated from blood samples of patients suffering from subacute bacterial endocarditis and sarcoidosis, respectively who received antibiotic treatment [19, 20]. Similar observations were made for patients with chronic brucellosis [21] and rheumatic fever caused by group A streptococci [22].

*Salmonella* was also reported to undergo L-form transition in the presence of cell wall-targeting drugs. Recent studies with *S*. Typhimurium showed that L-form can be readily induced by cefazolin [23]. These L form bacteria were also resistant to 3^rd^ generation cephalosporin [23]. Others envisaged L-form *Salmonella* to be responsible for the long term effects of attenuated bacterial vaccines, indicating their non-pathogenic nature [24]. On the contrary, several studies from 1970s suggested that L-form *Salmonella* was capable to cause disease. However, most of these publications were either not in English or unavailable in the databases.Development of L-form *Salmonella* in vivo and its role in pathogenesis require further elucidation

In this study, we induced transition to *S*. Typhi L-form in vitro by treating an ampicillin-sensitive strain (Ty2) with increasing concentrations of ampicillin over a period of 14 days. This exposure of a drug-sensitive *S*. Typhi strain to a gradient of ampicillin would mimic the *in vivo* condition during treatment of typhoid patients with antibiotics. We characterized the *S*. Typhi L-form and demonstrated its abilities to infect murine and human macrophage cell lines. Finally, we showed that L-form bacteria was capable to establish productive infection in a mouse model. This study, for the first time, explored in detail the potential pathogenic role of ampicillin-induced L-form of *S*. Typhi.

## MATERIALS AND METHODS

### Cells and reagents

Cell lines and cell culture reagents were procured from the American Type Culture Collection (ATCC) and Invitrogen, respectively. Bacterial culture media were purchased from BD Difco.

### Generation of *S*. Typhi L-form

Ampicillin sensitive *S*. Typhi Ty2 strain was sequentially exposed to increasing concentrations of ampicillin (0.05 ug/ml to 200 ug/ml) over a period of 14 days. Bacteria were allowed to grow for two days at each concentration of ampicillin before being transferred to media containing the next higher concentration of ampicillin.

### In vitro growth measurement

Growth curves of the parental and A_200_ strains were obtained by plotting optical densities of the bacterial culture media taken at 600nm wavelength (OD_600_) after different time intervals from the beginning of the culture.

### LPS extraction

LPS was purified by hot phenol method and was quantified by phenol-sulphuric acid method as described previously [25]. Briefly, bacteria were allowed to grow for 18 hrs in LB medium. The harvested cells were re-suspended in distilled water and lysed by sonication (7 Watt, 10 seconds pulse and 10 seconds interval for 20 mins). The cell lysates produced were treated with DNaseI (10 mg/ml) and RNase (5 mg/ml), followed by proteinase K (20 mg/ml) at 60°C for 1 hr. The lysate was then incubated with phenol at 65°C-70°C with vigorous shaking for 1 hr and the solution was allowed to cool down at 10°C before being centrifuged at 10000 rpm for10 min to separate the organic and aqueous phases. After that, double volume of ethanol was added to the aqueous phase and the solution was incubated overnight at −20°C for complete precipitation of the LPS.

### Peptidoglycan estimation

Peptidoglycan was measured according to the standard protocol [26]. One hundred microliter of bacterial culture was incubated with 50 µL of 1 M NaOH at 37 °C for 30 min, followed by the addition of 1 mL of concentrated sulphuric acid and incubation in a boiling water bath for 3 min. The mixture was cooled down rapidly on ice for 1 min and treated with 10 µL of 0.16 M Copper sulphate and 20 µL of 0.09 M p-hydroxy biphenyl at 30 °C for 30 min. Peptidoglycan was estimated by taking absorbance at 570 nm using a spectrophotometer.

### Cell culture

The human acute monocytic leukemia cell line, THP-1 was maintained in RPMI 1640 medium supplemented with 10% (vol/vol) heat-inactivated fetal bovine serum (FBS) and penicillin-streptomycin (50 µg/ml penicillin and 50µg/ml streptomycin). Cells were differentiated for 24 hr in the above culture medium, containing 100 nM phorbol 12-myristate 13-acetate (PMA). Experimentation with the differentiated THP-1 cells was done after an additional 24hr culture in absence of PMA. Mouse macrophage cell line RAW264.7 was cultured in Dulbecco’s modified Eagle medium (DMEM) supplemented with 10% FBS and penicillin-streptomycin.

### Gentamicin protection assay

This assay was done following the standard procedure [27]. Briefly, THP-1 cells were differentiated in 24-well culture plates (5 × 105 cells/well). RAW264.7 cells were similarly cultured in identical plates. Cells were infected at a multiplicity of infection (MOI) of 50 with *S*. Typhi, opsonized with the respective cell culture medium without antibiotics and were synchronized with the bacteria by centrifugation at 400 × g for 5 min. Infection was continued for 30 min at 37°C in the presence of 5% CO2. Extracellular bacteria were removed by repeated washing with 1× phosphate-buffered saline (PBS), followed by culturing the cells in media containing 100 μg/ml of gentamicin for 1 h and then, 15 μg/ml of gentamicin until the time of experiment. Cells were lysed with 1× PBS containing 0.25% Triton X-100, and intracellular CFU counts were carried out by overnight growth of the bacteria recovered from the cells on a Luria Agar, plate containing suitable antibiotics.

### Confocal microscopy

THP1 and RAW264.7 cells seeded on collagen-coated coverslips placed in 24 well plates (5× 10^5^ cells per well) were infected with *S*. Typhi. One day before the infection, the culture media were replaced by antibiotic-free media, containing dextran-conjugated Alexa fluor 594 (5mg/ml) (Thermo Fisher). After three washes with 1× PBS, coverslips were mounted on clean glass slides with fluorophore mounting media (Sigma). Dextran taken up by the cells emitted red fluorescence from the acidic vacuoles. Stained cells were viewed under a Zeiss LSM 710 confocal microscope, and bacteria were counted manually.

### In vivo experiments

All animal experiments were carried out following the protocols approved by the Institutional Animal Ethics Committee of ICMR-National Institute of Cholera and Enteric Diseases (NICED), Kolkata, India (Project license # PRO/103/May 2014-September 2017, Date 11.03.1999). ICMR-NICED adheres to the animal handling protocols, issued by the Committee for the Purpose of Control and Supervision of Experiments on Animals (CPCSEA), Ministry of Environment, Forest and Climate Change, Government of India.

An iron-overload mouse model was used for oral *S*. Typhi infection. Briefly, 6-to 8-weeks-old female BALB/c mice were injected intraperitoneally with desferrioxamine (Novartis, Switzerland; 0.025 mg/ gm of body weight) and Fe^3+^ (0.32 mg/ gm of body weight) 4 hr prior to the infection. Gastric acid was neutralized with 5% sodium bicarbonate 20 min before the oral infection with bacterial cultures, which were grown till OD_600_ reached around 1.0. Mice were infected orally with a sub lethal dose (5 × 10^5^ CFU) of *S*. Typhi and systemic infection was analyzed by the recovery of live bacteria from the liver, spleen, gall bladder, MLN and Peyer’s Patches on 2, 4 and 8 post-infection days. To this end, single-cell suspensions were prepared after the organs were lysed with 0.25% Triton X-100 (Sigma-Aldrich, USA) and plated on LA, containing suitable antibiotics.

For intra-peritonial infection, a sub lethal dose of *S*. Typhi (5 × 10^4^ CFU) was injected into the peritoneum of 6-to 8-weeks old BALB/c mice. Visceral organs (liver, spleen, and gall bladder) were collected at the same time points as above and CFU counts were done as stated before.

## Results

### *S*. Typhi L-form induction and characterization

Parental *S*. Typhi Ty2 strain was subjected to increasing concentrations of ampicillin over a period of 14 days. The bacterial strain obtained was called A_200_, since the dose escalation and subsequent culture was done in the presence of 200 ug/ml of Ampicillin. Visualization of the A_200_ strain under scanning electron microscope revealed gross morphological changes in the form of an elongated shape with higher cytoplasmic to nuclear volume (Fig. 1A and 1B). Each bacterial cell of the A_200_ strain contained more than one nucleoid region as compared to a single nucleoid in the parental strain (compare Fig. S1A and S1B), indicating compromised cell division in the former strain. Furthermore, comparison of the growth curves of the A_200_ and parental Ty2 strain showed significantly reduced growth for A_200_ strain (Fig. 1C). In contrast, naturally ampicillin-resistant *S*. Typhi strain, CT18 displayed a growth pattern similar to the parental Ty2 strain. However, when A_200_ was allowed to grow in the absence of ampicillin, it resumed the growth rate of the parental bacteria (Fig. S1C). In addition, most cells rapidly reverted back to the original morphology upon removal of ampicillin (compare Fig. 1 D and 1 E) Time kinetic studies revealed that majority of the A_200_ bacteria assumed the parental morphology with concomitant resumption of the growth rate to that of parental strain within a period of 4 hrs of culture in the absence of ampicillin (Fig. S1D). Together the above observations strongly suggested that A_200_ strain displayed the characteristics of the L-form bacteria. In agreement with the previous studies [28], L-form *S*. Typhi produced significantly reduced amounts of peptidoglycan (Fig. 1F) and LPS (Fig. S2A) than the parental strain. Similar to other L-form bacteria, A_200_ strain was more resistant to several antibiotics (Fig. S2B), further suggesting that it was indeed the L-form of *Salmonella* Typhi.

**Figure 1:**
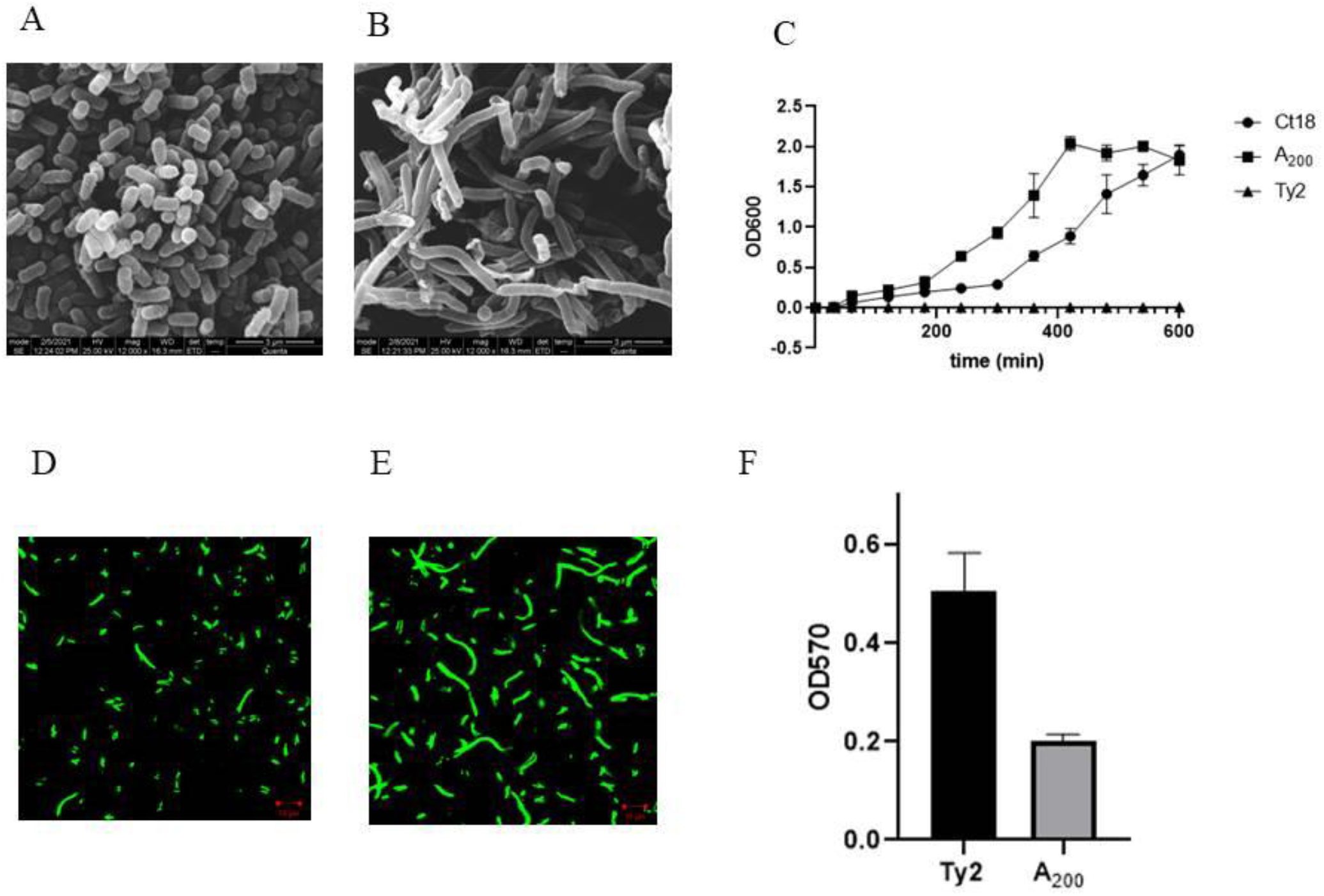
General characterization of A_200_: (A) Electron microscopy image of overnight grown WT Ty2. (B) Electron microscopy image of overnight grown A_200_. (C) Growth curve comparison of Ampicillin sensitive (Ty2), Ampicillin resistant (Ct18) and A_200_ L form. Data represents the mean of three independent experiments with SD value indicated by the error bar. (D) Quantification of total peptidoglycan in Ty2 and A_200_ strains. Data shown as mean of threeindependent experiments with SD value shown by error bar (E) Confocal microscopy of GFP containing A_200_ cells grown overnight in the absence of Ampicillin. (F) Confocal microscopy of GFP containing A_200_ cells grown overnight in the presence of Ampicillin.

### *S*. Typhi L-form was capable to infect macrophage cells

To compare infectivity of the parental Ty2 and the A_200_ strain, mouse macrophage cell line, RAW264.7 and human THP-1 cell-derived macrophages were infected with a multiplicity of infection (MOI) of 50, followed by gentamycin protection assay. At the 0 hr time point, higher number of intracellular A_200_ strain was recovered, indicating greater infectivity (Fig. S3 A and B). Twenty-four hours later, intracellular bacterial number increased for both the strains with A_200_ remaining more numerous, underscoring that the latter survived and multiplied within the mouse and human macrophage cell lines with equal efficiency as the parental Ty2 strain (Fig S3A and S3B). To gain further insights into the infectivity of A_200_ strain, confocal microscopy was performed. At both 0 hr and 24 hr time points, intracellular parental and A_200_ strains were co-localized with the acidic vacuoles (stained with Dextran-Alexa fluor 594) (Fig. 2, 3, 4, 5). Together the above results indicated that A_200_ strain efficiently infected macrophage cells and replicated within acidic vacuoles despite its altered morphology and slower growth *in vitro*. Given the higher infectivity of A_200_ strain, we investigated if it resulted in greater cell death. To this end, infected cells were stained with Annexin-V and propidium iodide (PI) and apoptosis was studied by flow cytometry. We observed that 65% of the Ty2-infected cells were apoptotic at 24 hrs post-infection, as compared with only 11% apoptosis for A_200_ infection. At 48 hrs post-infection, apoptosis further increased for Ty2- (∼70%), but not for A_200_-infected cells (Fig. 6 A - J). The results indicated that A_200_ strain induced minimal cell death despite more efficient infection *in vitro*.

**Figure 2.**
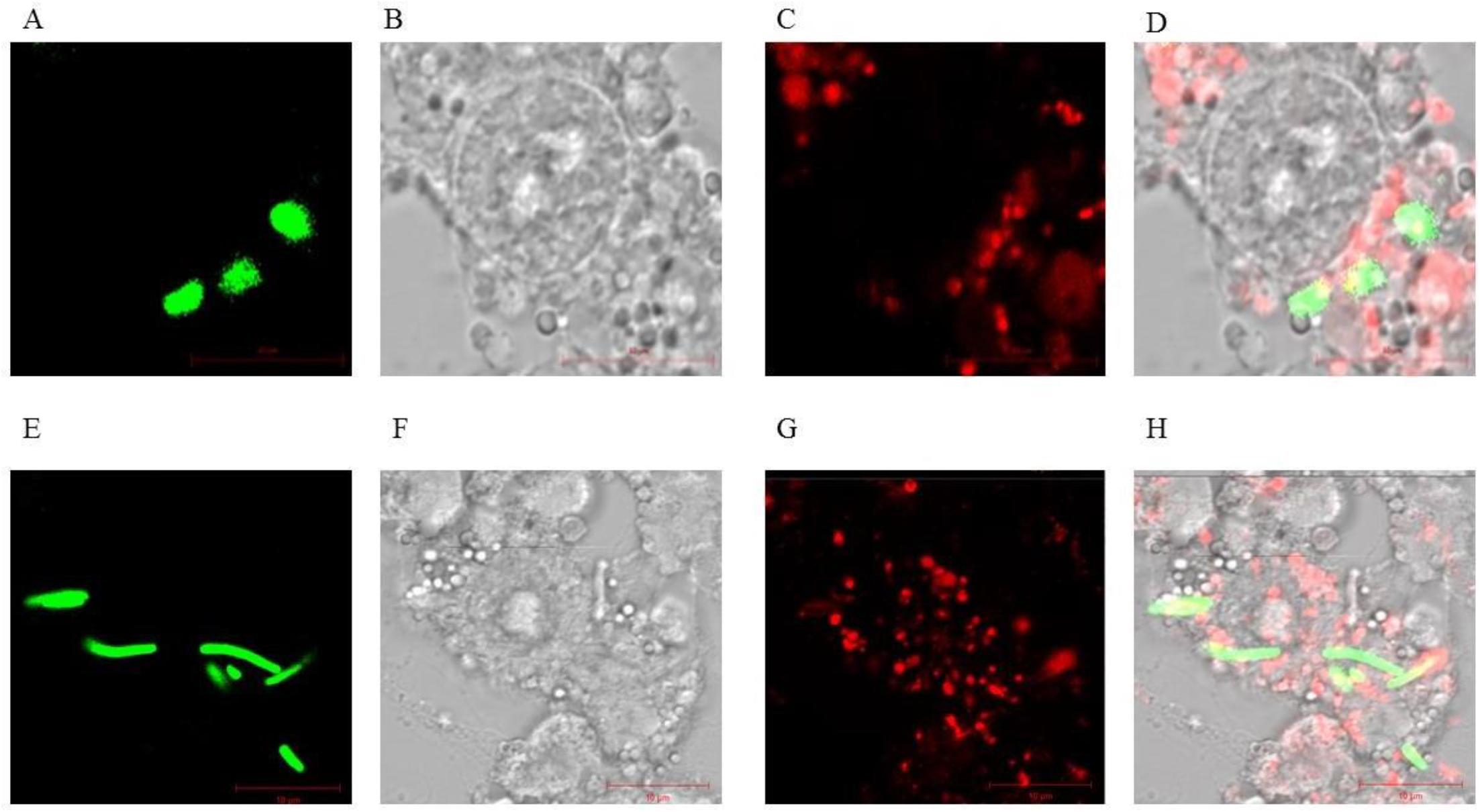
Infection of murine RAW cell line with GFP- Ty2 and GFP- A_200_ L-form at 0 hr: Representative confocal microscopy images showing intracellular Ty2 (upper panel) and A_200_ (lower panel) bacteria at 0hr post infection. (A and E) GFP containing bacteria. (B and F)DIC image of RAW cells. (C and G) Acidic vacuoles stained with Dextran Alexa Fluor. (D and H) Merge of all the above filters.

**Figure 3:**
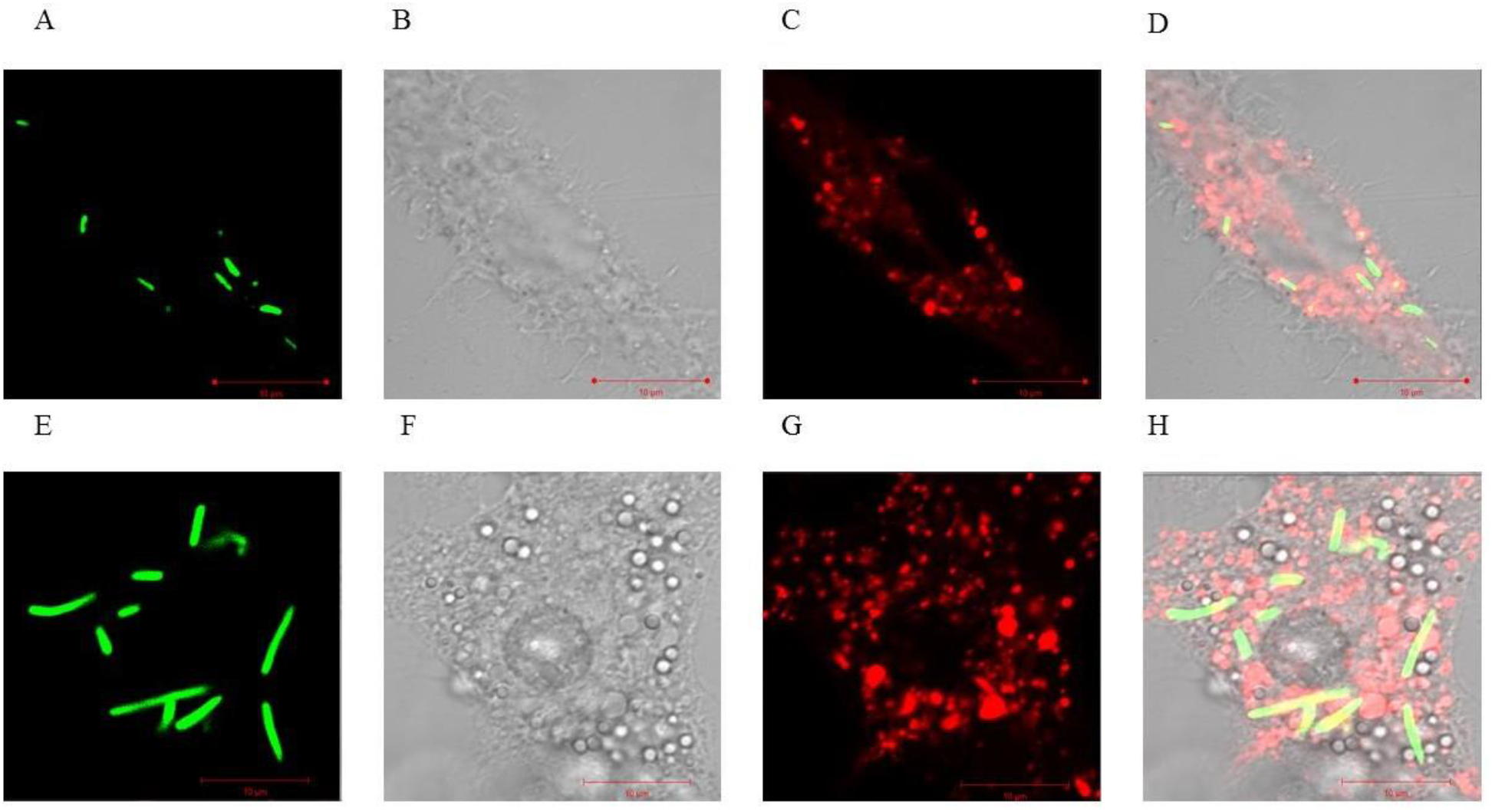
Infection of murine RAW cell line with GFP- Ty2 and GFP- A_200_ L-form at 24hr: Representative confocal microscopy images showing intracellular Ty2 (upper panel) and A_200_ (lower panel) bacteria at 24hr post infection. (A and E) GFP containing bacteria. (B and F) DIC image of RAW cells. (C and G) Acidic vacuoles stained with Dextran Alexa Fluor. (Dand H) Merge of all the above filters.

**Figure 4:**
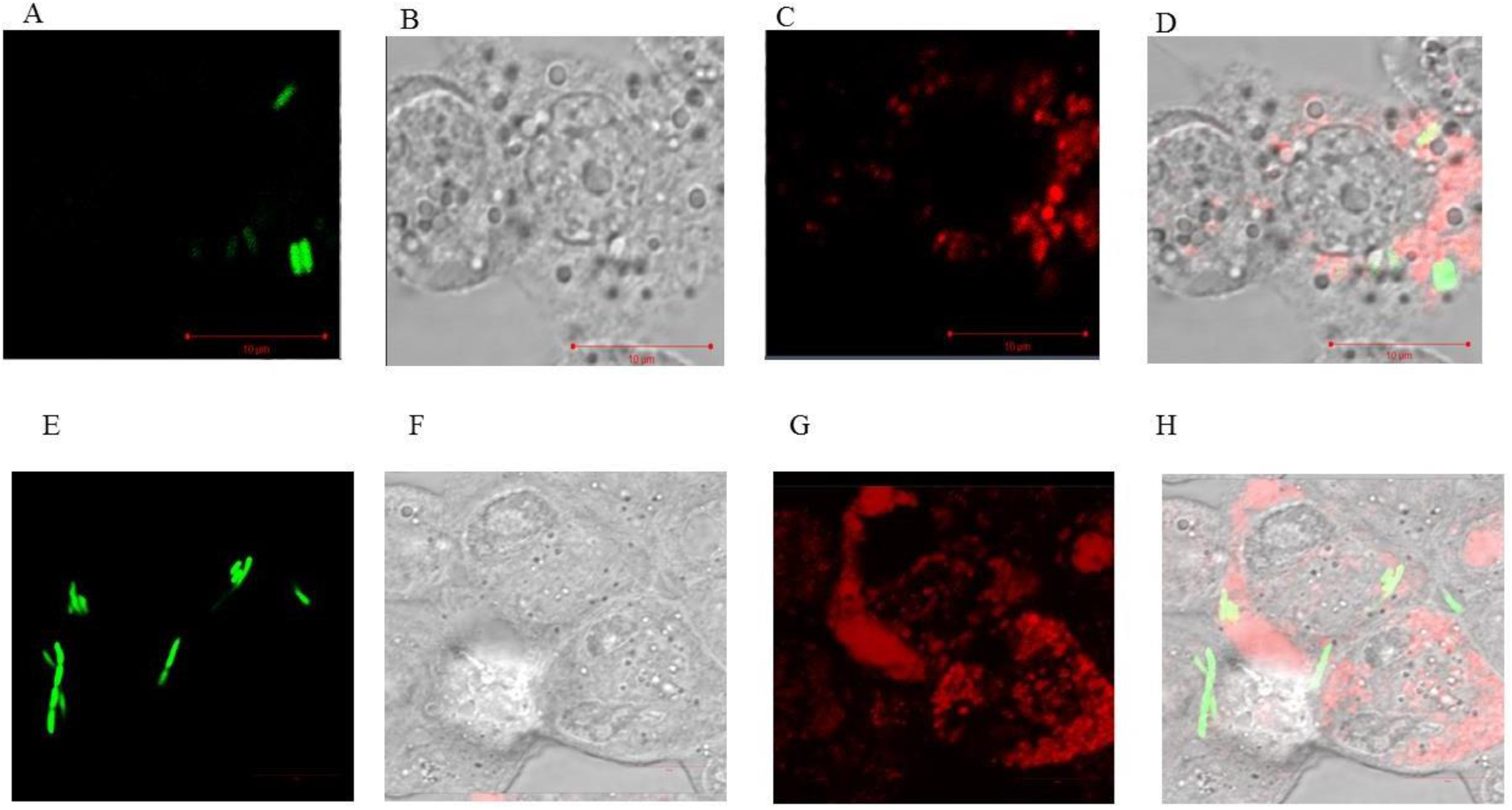
Infection of THP-1 cell line with GFP- Ty2 and GFP- A_200_ L-form at 0 hr: Representative confocal microscopy images showing intracellular Ty2 (upper panel) and A_200_ (lower panel) bacteria at 0hr post infection. (A and E) GFP containing bacteria. (B and F) DICimage of THP-1 cells. (C and G) Acidic vacuoles stained with Dextran Alexa Fluor. (D and H)Merge of all the above filters.

**Figure 5:**
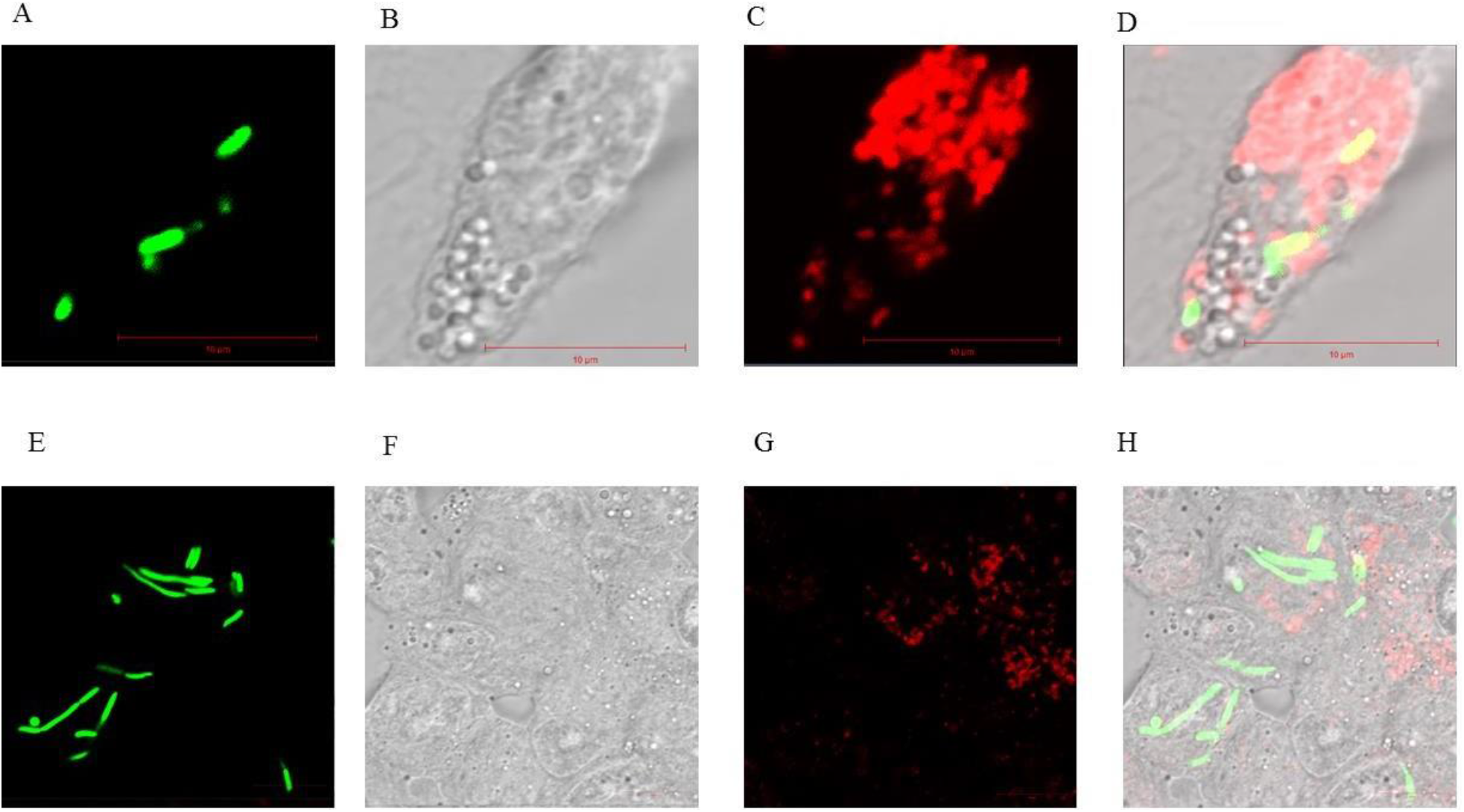
Infection of THP-1 cell line with GFP- Ty2 and GFP- A_200_ L-form at 24 hr: Representative confocal microscopy images showing intracellular Ty2 (upper panel) and A_200_ (lower panel) bacteria at 24 hr post infection. (A and E) GFP containing bacteria. (B and F) DIC image of THP-1 cells. (C and G) Acidic vacuoles stained with Dextran Alexa Fluor. (D and H) Merge of all the above filters.

**Figure 6:**
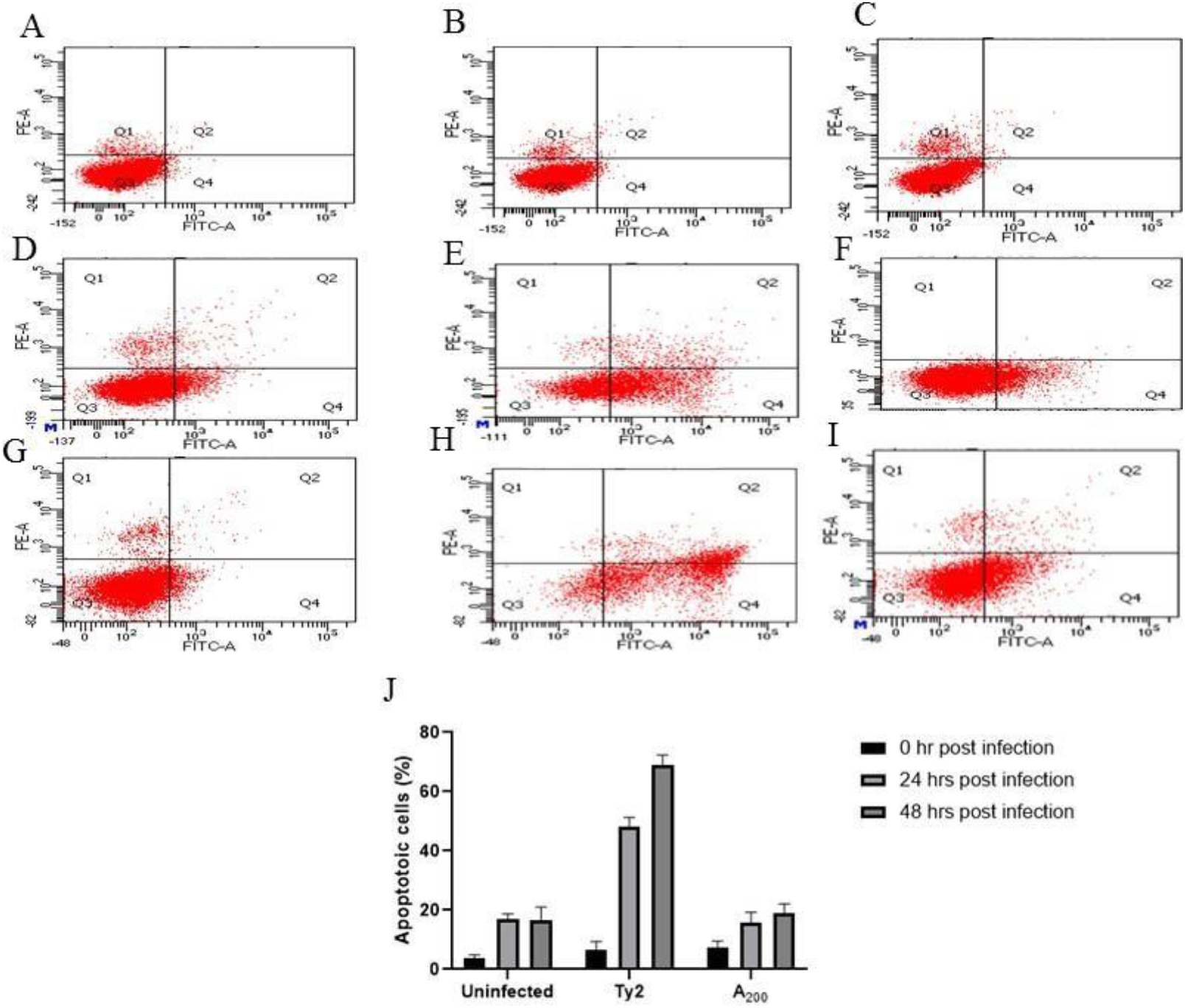
Apoptosis induced by Ty2 and A_200_ in THP-1 cell line: Representative flow diagram of Ty2 and A_200_ infected THP-1 cells. (A) uninfected THP-1 cells, 0 hr. (B) Ty2 infectedTHP-1 cells, 0 hr. (C) A_200_ infected THP-1 cells, 0 hr. (D) uninfected THP1 cells, 24 hr, (E)Ty2 infected THP-1 cells, 24 hr. (F) A_200_ infected THP-1 cells, 24 hr. (G) Uninfected THP1cells, 48 hr (H) Ty2 infected THP1 cells, 48 hr (I) A_200_ infected THP1 cells, 48hr. (J) Quantification of apoptotic cell by flow cytometry after annexin V/ PI staining of treated and untreated Thp-1 cell lines after 0 hr, 24 hrs and 48 hrs post infection. Data represents the meanof three independent experiments with SD value indicated by the error bar.

### *S*. Typhi L-form established infection in mouse model

The ability of A_200_ strain to infect macrophage cell lines prompted us to investigate if the strain was also capable to establish infection *in vivo*. Mice were infected orally as well as through the intraperitoneal (IP) route with a sub-lethal dose of the parental Ty2 or A_200_ strain. Bacterial load was measured at 2, 4 and 8 days post-infection (dpi) in the visceral organs. Equal and high number of both the bacterial strains were recovered from the liver, spleen and gall bladder after 2 days of intraperitoneal infection, indicating that A_200_ was equally competent as the parental strain to infect live animals (Fig. 7). Bacterial load further increased for both strains at 4 dpi, underscoring similar efficiency of survival and replication *in vivo*. However, there was a drastic reduction of load in the visceral organs at 8 dpi that suggested pathogen clearance by the host immune system. Interestingly, the visceral load for A_200_ was significantly higher compared with the parental strain at the later time point. This observation indicated that A_200_ strain was perhaps more competent in evading the host immune system. As opposed to intraperitoneal infection, substantial bacterial recovery after oral administration was only observed at 4 dpi, suggesting delay in systemic invasion following intestinal colonization (Fig 8). Moreover, significantly fewer A_200_ bacteria than the parental strain were recovered from the internal organs at this time point. Given that both the strains survived and replicated in the visceral organs with equal competence and A_200_ strain was more efficient to resist clearance by the host (Fig 7), the above result suggested greater systemic invasion by parental *S*. Typhi from the intestine. However, greater resistance of A_200_ strain to host clearance led to minimization of the difference between the bacterial loads of the two strains in the internal organs at 8 dpi (Fig. 8).Together, the above results suggested that the growth of *S*. Typhi L-form was similar to the parental bacteria after oral and intraperitoneal infection, in contrast to its slower growth in thein vitro cultures. However, L-form was less efficient in systemic translocation from the gut, but resisted clearance by the host more efficiently.

**Figure 7:**
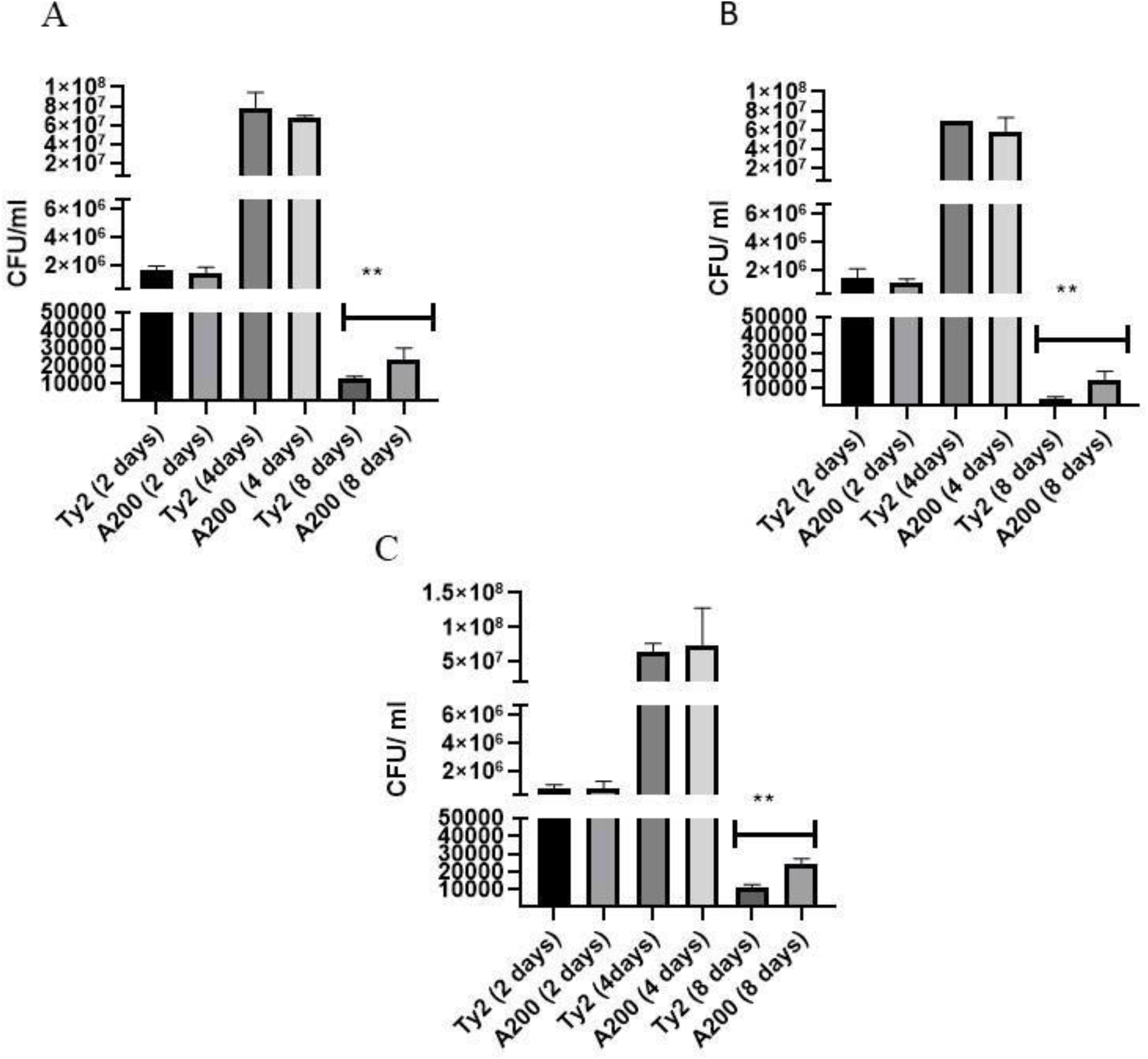
Infection of mice model by A_200_ L form (IP): Mice (n = 5) were infected intraperitoneally with A_200_ L-form. Bacteria was isolated from different organs, (A) liver, (Bspleen, (C) Gall bladder after 2, 4, and 8 dpi and platted on LA plates. CFU was calculated and plotted. Data represents the mean of three independent experiments with SD value indicatedby the error bar.

**Figure 8:**
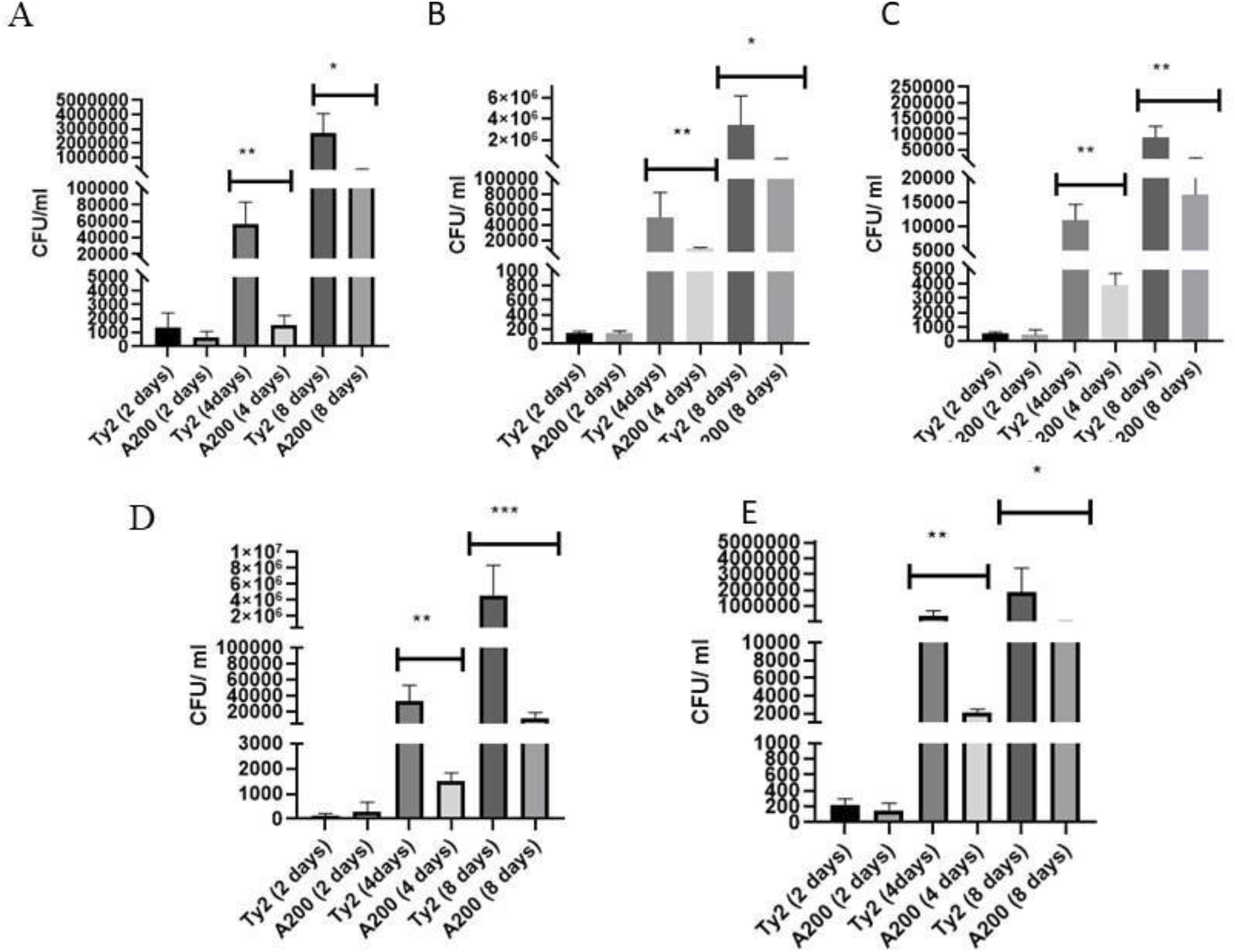
Infection of mice model by A_200_ L form (oral): Mice (n = 5) were infected orally with A_200_ L-form. Bacteria was isolated from different organs, (A) liver, (B) Spleen, (C) Gall bladder, (D) MLN, (E) Payers’ patch after 2, 4, and 8 dpi and platted on LA plates. CFU was calculated and plotted. Data represents the mean of three independent experiments with SD value indicated by the error bar.

## Discussion

Antibiotic treatment cures bacterial infection either by killing the pathogen or halting their growth, leading to clearance of bacteria by the host immune system. However, antibiotic concentration in the tissues does not reach the optimum level after the first dose of the drug, rather it increases with time till it reaches the desired concentration. Thus, bacterial population within tissues is subjected to a progressively increasing gradient of antibiotic. In this study, we aimed to investigate the effects of an increasing concentration gradient of the antibiotic drug Ampicillin on *Salmonella* Typhi. Ampicillin inhibits transpeptidases and kills bacteria by blocking cell-wall synthesis. When ampicillin-sensitive *S*. Typhi was subjected to progressively higher doses of the drug over a period of 14 days, the bacteria changed the morphology, shut down cell wall and LPS synthesis and was able to grow in the presence of 200 ug/ml of ampicillin. This phenomenon was not observed when *S*. Typhi Ty2 strain was cultured in the presence of 200 ug/ml ampicillin from the beginning that killed all the bacterial cells. This contrasts with *Salmonella* Typhimurium, where L-form was induced by culturing antibiotic-sensitive bacteria with another cell wall inhibiting drug called cefazolin at a much lower dose than the MIC from the beginning [23]. Increasing concentration gradient of ampicillin led to the transition of *S*. Typhi Ty2 strain to an L-form bacteria, as characterized by drastically reduced cell wall and LPS production, an elongated shape with higher cytoplasmic volume and more than one nucleoid per cell, and significantly compromised growth in the *in vitro* cultures. However, characteristics of the parental strain were largely restored when ampicillin was removed from the culture media, indicating that the changes were reversible. The morphological changes observed in this study were similar to that observed for cefazolin-induced L-form of *S*. Typhimurium [23]. Similar phenotypic changes were reported by studies with MDR *E. coli* A9 strain subjected to 20X MIC of ampicillin [29].

L-form of several pathogenic species were isolated from different organs of individuals suffering from symptomatic infections as well as the carriers of the disease [30]. Isolation of the L-forms from the carriers particularly fascinated the microbiologists for several decades about the actual role of this form in the pathogenesis of a disease. *Streptococcus pyogenes* L-form was found to infect an animal model and was shown to persist over a longer period of time compared with the walled form. This L-form bacterium was phagocytosed by macrophages and survived and replicated inside these cells [31]. On the contrary, L-form of another gram positive bacterium, *Corynebacterium diphtheriae* failed to cause productive infection and disease in the animal model. The authors concluded that defects in the cell wall formation abrogated diphtheria toxin production, leading to loss of infectivity [32]. Unlike the Gram positive bacteria, pathogenesis of Gram negative bacterial L-forms remains largely unexplored.

In this study, *S*. Typhi L-form was shown to infect murine as well as human macrophage cell lines. Gentamycin assay and confocal microscopic studies revealed active multiplication of A_200_ strain inside these cells. Moreover, infectivity of A_200_ strain was higher compared with parental *S*. Typhi. This may be explained by reduced LPS synthesis by the A_200_ strain, since LPS was earlier reported to inhibit phagocytosis by macrophages. However, higher infectivity of the L-form did not result in greater cell death, suggesting the potential for longer intracellular persistence.

A_200_ strain was equally efficient as the parental *S*. Typhi strain to establish infection in a mouse model. When administered IP, equivalent number of the bacterial strains was isolated from different organs at 2 and 4 dpi, indicating that both the strains infected and multiplied with equal efficiently *in vivo*. However, despite significant reduction in bacterial load at 8 dpi, suggesting bacterial clearance by the host immune system, the load for A_200_ strain in the host tissues remained considerably higher compared with the parental bacteria. This suggested that A_200_ was more efficient in evading the host immune system. This is surprising given that cell wall confers protection to bacteria against the host machinery. However, similar results were observed for *Streptococcus pyogenes* infection, where the L-form persisted for a longer period in the tissues and was thought to be responsible for recurrent infection by the bacteria [31]. Perhaps, absence of cell wall helped the L-form bacteria to avoid activation of the host immune system, thereby evading a clearance response by the host.

Surprisingly, A_200_ displayed different behavior inside the host upon oral delivery when compared with IP administration. Small, but equal numbers of the parental and A_200_ strains were isolated from different organs of mice at 2 dpi, indicating equally efficient gut colonization and early translocation to the systemic sites. However, at 4 dpi, recovery of A_200_ strain was less than the parental bacteria. This was not due to slower replication of A_200_ strain, as proved by the results under Fig 7. This was not due to more rapid clearance of *S*. Typhi L-form from the systemic circulation either, since both the strains were isolated at higher numbers at 8 dpi than 4 dpi. Instead, cell-wall deficient A_200_ strain could have activated the intestinal immune system to a greater magnitude than the parental strain, thus preventing systemic translocation from the intestine at later time points. The exact mechanisms underlying different relative recovery of the parental and L-form *S*. Typhi after oral infection requires further investigation.

*S*. Typhi was earlier shown to spontaneously convert to L-form in the gall bladder [33]. In our study, A_200_ strain was able to colonize the gall bladder after both oral and intraperitoneal infections. Gall bladder colonization and faecal shedding are the most important factors behind *S*. Typhi transmission from otherwise healthy carriers [34]. The prevailing concept is that *S*. Typhi shed in the stool needs to revert back to the walled phenotype to become infective for a new host. Our study showed that *S*. Typhi L-form was equally capable to cause infection, which suggests more potent human to human transmission.

## Conclusions

No studies on the pathogenesis of the L-forms *Salmonella spp* are currently available. We, for the first time, studied pathogenesis of *Salmonella* Typhi L-form after oral as well as intraperitoneal infection. This study revealed that exposure of *Salmonella* Typhi to an increasing concentration gradient of ampicillin during the treatment of human infection fails to clear the entire bacterial population from the host. Rather, it may render *Salmonella* resistant to the antibiotic without notable genetic mutations. In this state, cell-wall deficient and elongated bacteria infects macrophages and multiply efficiently *in vivo*.

## Data availability

All data supporting the final conclusions are available in figures

